# Auditory Decisions in the Supplementary Motor Area

**DOI:** 10.1101/2020.10.20.347864

**Authors:** Isaac Morán, Javier Perez-Orive, Jonathan Melchor, Tonatiuh Figueroa, Luis Lemus

## Abstract

In human speech and communication across various species, recognizing and categorizing sounds is fundamental for the selection of appropriate behaviors. But how does the brain decide which action to perform based on sounds? We explored whether the premotor supplementary motor area (SMA), responsible for linking sensory information to motor programs, also accounts for auditory-driven decision making. To this end, we trained two rhesus monkeys to discriminate between numerous naturalistic sounds and words learned as target (T) or non-target (nT) categories. We demonstrated that the neural population is organized differently during the auditory and the movement periods of the task, implying that it is performing different computations in each period. We found that SMA neurons perform acoustic-decision-related computations that transition from auditory to movement representations in this task. Our results suggest that the SMA integrates sensory information while listening to auditory stimuli in order to form categorical signals that drive behavior.

## Introduction

Recognition and categorization of sensory information are essential for deciding which action to take (Parker & Newsome, 1998; Freedman et al., 2001; Romo et al., 2003; Seger & Miller, 2010). In particular, categorizing dynamic sounds into classes is necessary for social communication in various species (Prather et al., 2009; Miller et al., 2003), including monkeys (May, Moody & Stebbins, 1989) and humans, where grouping of discrete sounds into words is fundamental for speech recognition (Repp, 1984; Leonard & Chang, 2014). However, it is not clear how the brain links the recognition of complex sounds to behavior. In primates, auditory information is carried from the auditory cortex to the prefrontal and premotor cortices via ventral and dorsal auditory streams (Romanski et al., 1999; Kusmierek & Rauschecker, 2014). Recent studies with macaques have shown that premotor regions may represent an embodiment of acoustic recognitions (Archakov et al., 2020). The premotor supplementary motor area (SMA) has been linked to voluntary action and cognitive control of movement (Nachev et al., 2008; Lara et al., 2018; Shima & Tanji, 2000). The SMA also participates in working memory and decision making during tactile discrimination (Romo et al., 1997; Hernandez et al., 2002; Lemus et al., 2007), imagery, and it links auditory information with motor programs (Lima et al., 2016; Vergara et al., 2016). Similarly, in songbirds, the premotor HVC nucleus is involved in coding complex sounds and orchestrates singing (Prather et al., 2009).

Nevertheless, few studies have been published on naturalistic sound coding (Rauschecker, 1998; Gentner & Margoliash, 2003; Tsunada et al., 2011; Town et al., 2018), and even fewer in behaving non-human primates (Ng et al., 2009; Scott et al., 2012; Chandrasekaran et al., 2013; Melchor et al., 2020). This could be because there is still plenty to learn from experiments using simple, non-naturalistic sounds that do not require behavior (e.g. pure tones, amplitude-modulated noise; Liang et al., 2002; Selezneva et al., 2006; Bendor & Wang, 2008; Yin et al., 2020). Additionally, although macaques have brain organizations that are similar to those of humans and although they, too, use vocalizations to communicate (Chang et al., 2010; Daube et al., 2019), some consider that non-human primates are challenging for training and studying auditory behavior (Ng et al., 2009; Scott et al., 2012). However, Melchor et al. (2020) recently showed that macaques attend to and memorize numerous naturalistic sounds. Therefore, in macaques, the SMA is amenable for studying the neural correlates of decisions within an auditory categorization task.

Here we present data from extracellular recordings of SMA neurons from two rhesus monkeys trained to discriminate between two groups, or categories, of naturalistic sounds and then choose an action. We also tested acoustic morphings to reveal the underlying responses to linear physical changes rather than discrete categorizations. We found robust categorical responses at both the single neuron and population levels. A small number of neurons were also found to carry information about individual sensory stimuli during the auditory period. By mapping the neural population activity during different periods of the task, we found that it organizes into orthogonal subspaces during the task’s auditory and movement periods. Finally, the linear morphings of one stimulus into another indicate that the coding in SMA changes from sensory, during the auditory period, to categorical during the movement period. Taken together, our results support the hypothesis that SMA integrates auditory information necessary for a categorization that provides action signals for behavioral choices.

## Results

### Monkeys discriminate between learned naturalistic sounds

We trained two monkeys (*Macaca mulatta*) to perform an auditory recognition task. During the task, the monkeys obtained a drop of liquid as a reward for releasing the lever only after identifying a target sound (T). A T followed 0, 1, or 2 non-target sounds (nT). In other words, the T could appear in three different temporal positions (P1, P2, and P3; Figs. 1a,c, see Methods). It is important to note that we incorporated a delay and a green visual cue (VC) after each sound to disentangle auditory and movement signals. Thus, after a T, the monkeys waited for a 0.5 s delay before a 0.5 s VC-release period (Figs. 1a,c). The monkeys learned, by trial and error, to recognize up to fourteen sounds (half of them T, half nT) belonging to four different classes: conspecific vocalizations, interspecific vocalizations, words, and artificial sounds (Figs. 1b,d; see Methods). Once the monkeys performed above 85% correct responses (Fig. 1d), we recorded single neurons (1341 from monkey 1; 524 from monkey 2) from SMA (Fig. 1c (inset) during task execution).

**Figure 1.**
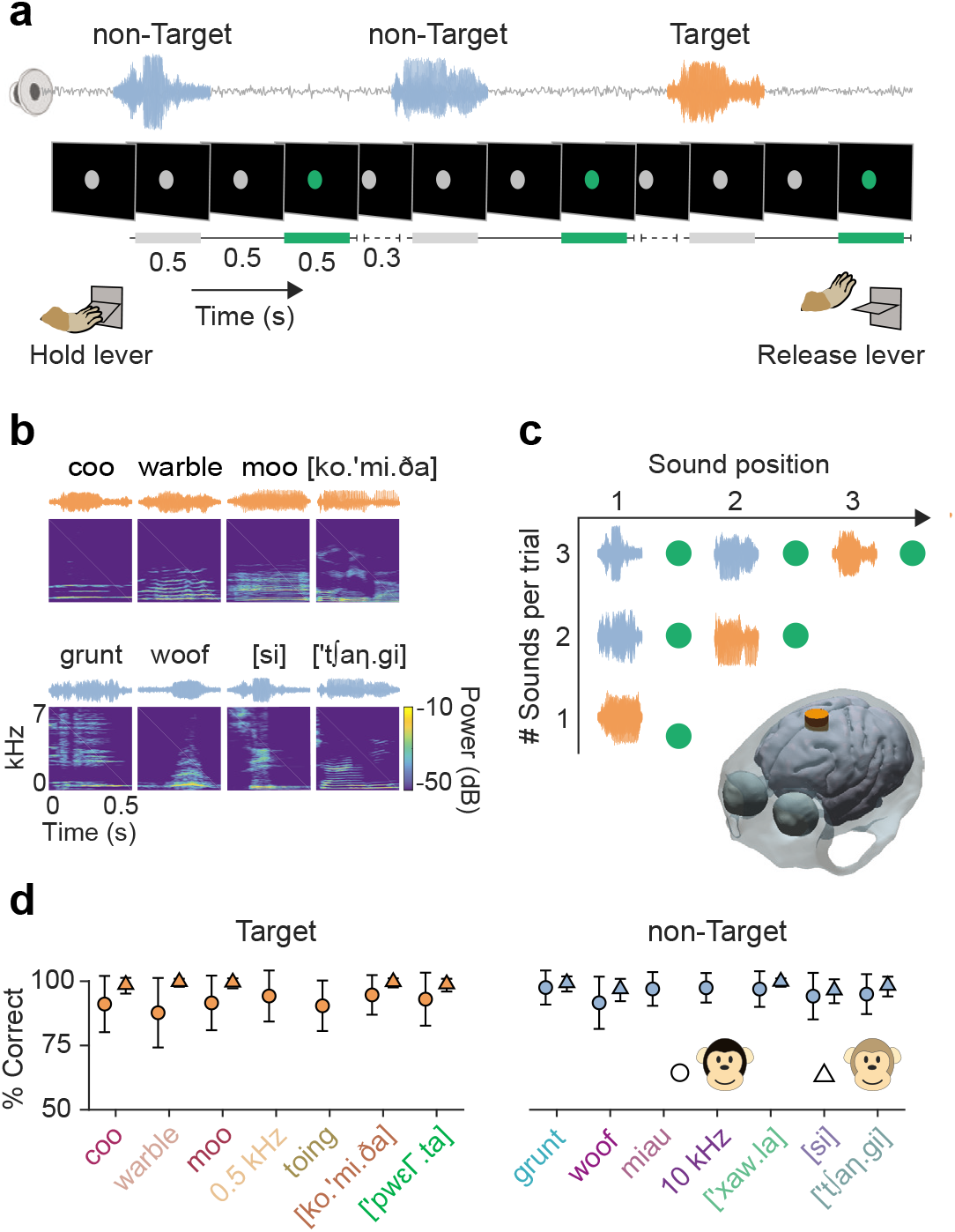
Acoustic recognition task and behavior. **a)** Sequence of events in a trial: The monkey pressed the lever after a gray visual circle indicated the beginning of a trial. Then, a playback of one to three sequential sounds commenced. Each sound lasted 0.5 s followed by a 0.5 s delay and a 0.5 s green visual circle (go cue). The monkey received a reward for releasing the lever within a 0.7 s window after the green cue following a Target sound (T) (red sonograms). False alarms occurred when the monkey released the lever earlier, e.g. during a non-Target sound (nT) (blue sonograms). Finally, not releasing the lever during the T response window counted as a miss trial. **b)** Examples of sonograms and spectrograms of four T (top) and four nT (bottom) sounds. T and nT included four classes: conspecific vocalizations, interspecific vocalizations, words, and artificial sounds. **c)** Types of trials regarding the number of sounds. **Inset,** Brain scheme of MRI of monkey 2 showing the location of the recording chamber over SMA. **d)** Percentage of correct responses for T (red) and nT (blue) for each monkey (monkey 1: circles; monkey 2: triangles). The color code of stimuli will be referenced in the following figures.

### Single units and population activity represent naturalistic sounds

Rasters of a characteristic SMA neuron (Fig. 2a) show a substantial firing rate change during a T regardless of its position (i.e. P1, P2, or P3). In other words, the responses show a clear distinction between T and nT, but with small or no differences within each category (Figs. 2b and c). The majority of neurons recorded in both monkeys (68.5% and 85.9%, respectively) exhibited this type of categorical coding (Fig. 2d), as measured with the f-statistic and bootstrapping (see Methods). Interestingly, a small percentage of neurons displayed significant coding for specific sensory stimuli within the T and nT categories (2.4% and 1.1% for T; 1.1% and 2.1% for nT in each monkey, respectively; Fig. 2d). As shown in Figures 2b, 2c, and 2e, neurons with this particular sensory coding did this during the auditory periods when sounds were presented (0.0-0.5 s). In contrast, the f-statistic shows that the categorical coding starting at the end of auditory periods rises further during the delays (0.5-1.0 s) and movement periods (1.0-1.5 s; Fig. 2e).

**Figure 2.**
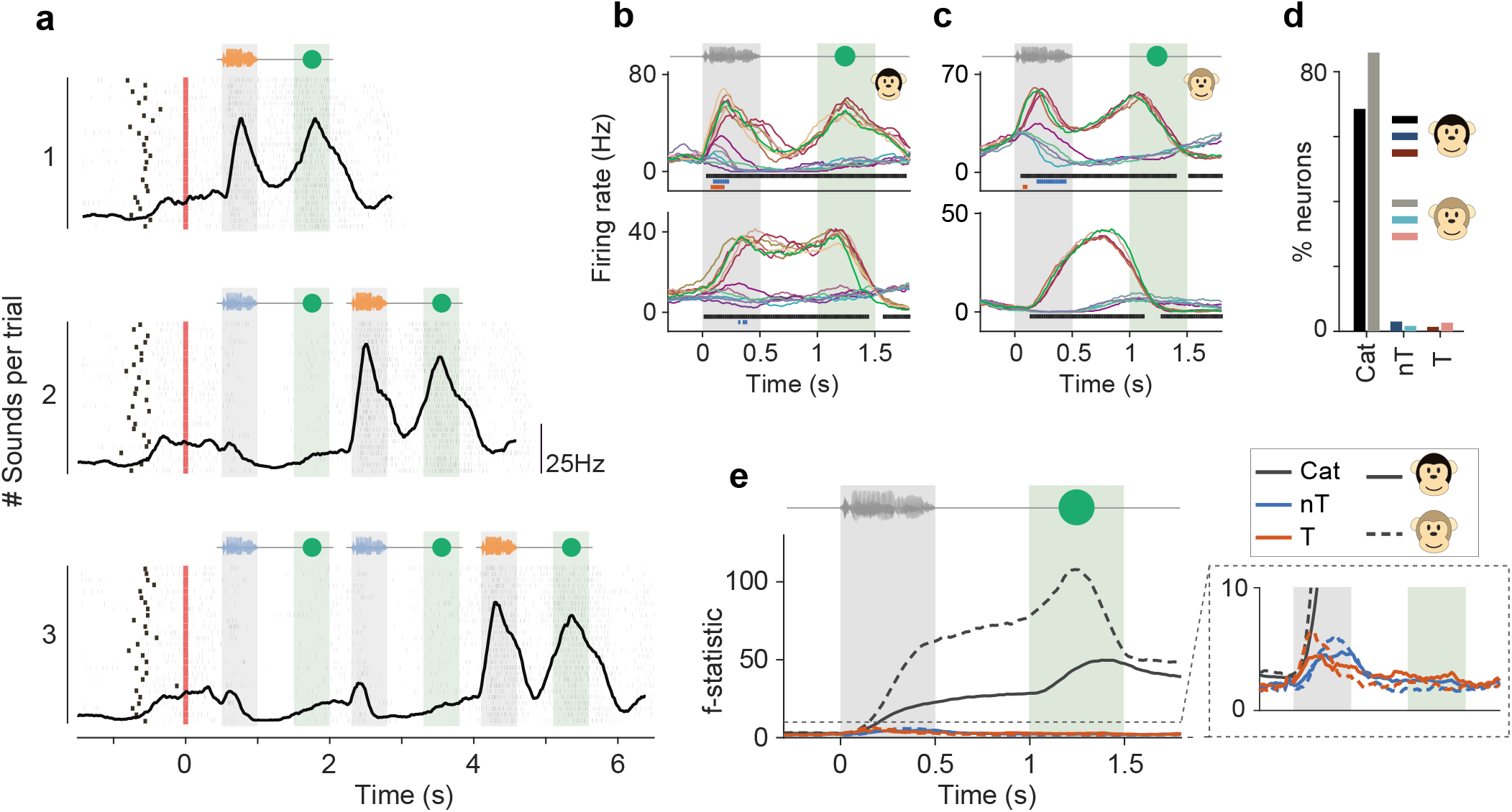
Individual SMA neurons code behavioral categories. **a)** Rasters and peristimulus time histograms (PSTHs) of a representative neuron from monkey 1. Each row of ticks is one trial aligned with the lever press; each tick is an action potential. Each plot corresponds to the possible number of sounds in a trial. Thick black ticks indicate the beginning of a trial. Red ticks, lever pressings. **b)** Example PSTHs of two neurons for monkey 1 aligned with the start of the sounds. The color lines represent the PSTHs for each T and nT stimulus. The lines below (black, categorical; red, T; blue, nT) indicate bins of significant coding. T stimuli are depicted in red and nT in blue palettes, as in Figure 1d. The top neuron is the same as in (a). **c)** Same as (b) but for monkey 2. **d)** Percentage of recorded SMA neurons with statistically significant coding of behavioral categories (Cat) or sensory information (nT and T), and for each monkey. **e)** Average F-statistic for selective SMA neurons as a function of time for both monkeys (monkeys 1: continuous lines; 2: dashed lines); **Inset**, a magnification showing small F-statistic values for the few neurons with significant sensory coding for particular T or nT sounds.

To study the neuronal population dynamics, we used principal component analysis (PCA), a dimensionality reduction technique. Similarly to the observed results of individual neurons, the population dynamics differentiated between T and nT, independently of the position in which T appears (Fig. 3a). The component explaining most of the variance shows a small increase at the beginning of the T stimulus, followed by a larger increase during the movement period, as expected for this premotor area. The second component shows acoustic-anticipatory activity that continues to rise with Ts but decreases for nTs, providing a clear auditory-based signal for premotor activity. The third principal component shows a large increase at the beginning of the trial and an additional increase before the lever release indicating the end of the trial.

**Figure 3.**
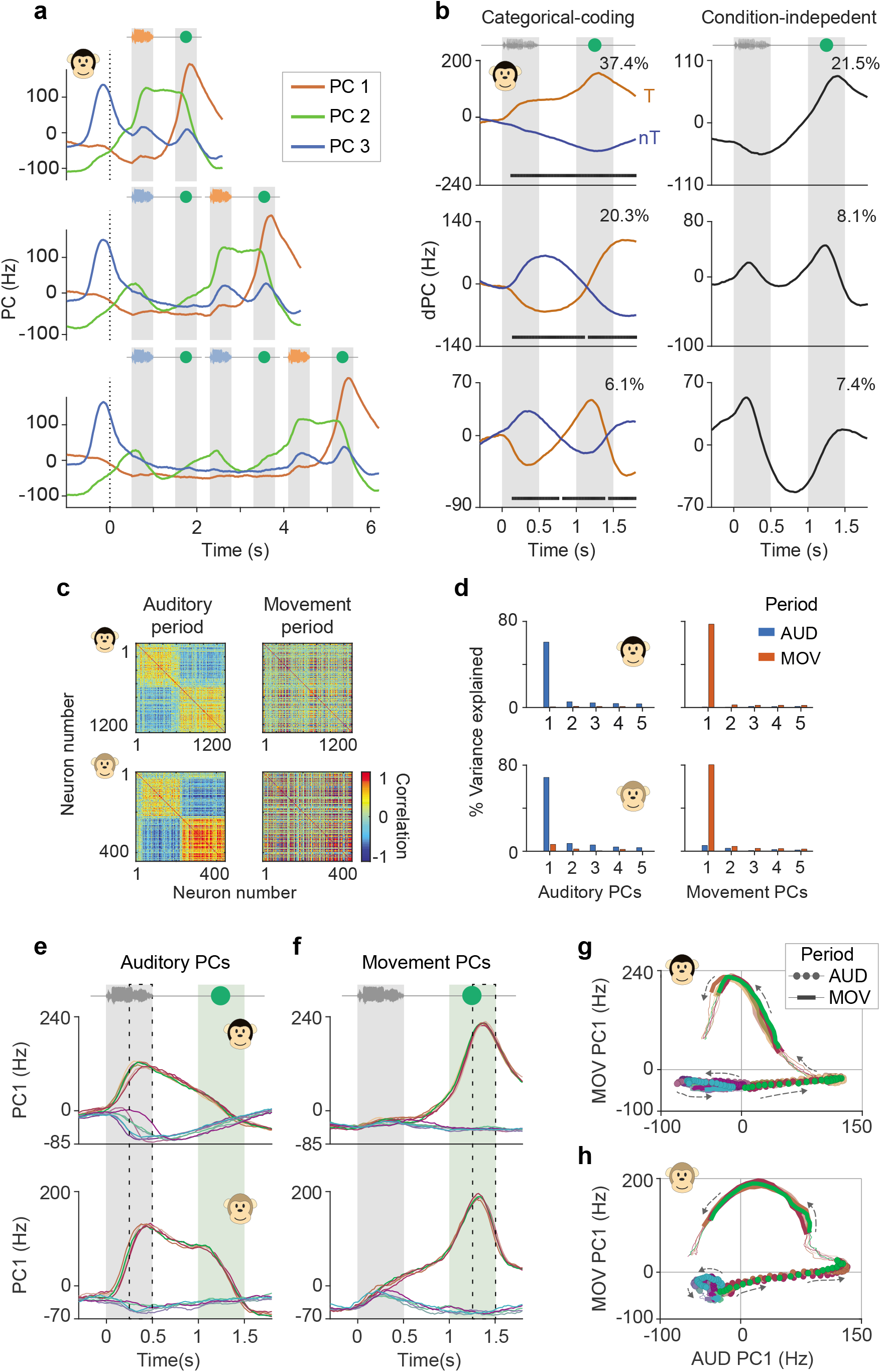
Categorical responses of SMA population project in auditory and movement orthogonal subspaces. **a)** Projections of the neural population responses in the first three principal components. The plot is grouped by the number of sounds, as in Figure 2a. Notice PC1 codes for lever release, PC2 for anticipation of sounds together with the likelihood of a T, and PC3 for Ts and movements. **b)** Left: First three dPCs for categorical coding. Each trace corresponds to T (red) and nT (blue) projections. Right: First three condition-independent dPCs. The numbers on the top right are the percentages of explained variance of all dPCs. The first categorical coding components explain a greater amount of variance compared to the condition-independent components. **c)** Correlation matrices for auditory (left) and movement (right) periods of all neurons of monkeys 1 (top) and 2 (bottom). Each dot corresponds to the correlation between two neurons. Matrices for both periods sorted as a function of the auditory period. The groups of correlated neurons that exist in the auditory period disappear in the movement period. **d)** Percentage of explained variance by each of the top five acoustic (left) and movement (right) PCs projected onto the population activity during auditory and movement periods. **e)** Projections of the neural population activity onto the first PC of the auditory subspace for monkeys 1 (top) and 2 (bottom). **f)** Same as (e), but for the movement subspace. **g)** Two-dimensional projection of the auditory and movement orthogonal subspaces for monkey 1. Circles and thick lines show the auditory and movement periods, respectively. Arrows indicate flow of time. Same color code as in Figure 1d. **h)** Same as (g), but for monkey 2.

We also used demixed PCA (dPCA), a dimensionality reduction tool that can separate the effects on the population response of different task parameters (Kobak et al., 2016), to separate the effect of the auditory stimulus coding from the condition-independent activity (Fig. 3b). Since the previous results from single units showed strong categorical codification, we computed dPCA for the T and nT categories. The demixed principal components (dPCs) with the most weight explaining the variance of the population activity are those for the categorical coding (first two categorical coding dPCs explain 57.7% and 65.3% of the total variance for each monkey, respectively, while the first two condition-independent dPCs explain 29.6% and 29.5% of the variance). For both monkeys, the dPCs separated the T and nT trajectories during the auditory periods. For dPC1, this separation persisted during the delay and reached its maximum during the movement period (Fig. 3b, top). Here, dPC2 and dPC3 shift their signs in the delay or during the movement period. The neural activity projections over the condition-independent dPCs are associated mainly with the task’s temporal dynamics.

### Auditory and movement computations occupy different subspaces

After observing the differences between auditory and movement period responses of single neurons and the SMA population, we set out to determine whether such representations proceeded from distinct population organizations, each coming together at different time periods to perform different computations (Elsayed et al., 2016). Figure 3c highlights the correlation matrices for the full neural population, organized so the structure during the auditory period is highlighted, while keeping this same ordering for the movement period. The correlation matrices show two large groups of neurons for both monkeys, with high correlations during the auditory period, but not during the movement period (Fig. 3c). To evaluate whether this difference was due to two different populations, we calculated a simple epoch-preference index (Elsayed et al., 2016; see Methods). If the preference index distribution for all neurons is bimodal, this implies two pools of neurons. However, we found evidence that both monkeys’ data came from a unimodal distribution (Hartigan’s dip test p= 0.98 and p = 0.96 for monkeys 1 and 2, respectively), suggesting that the auditory and movement activities did not come from two different populations.

We used PCA to investigate whether these population responses during the auditory and movement epochs occupy partially overlapping or orthogonal subspaces within the neural activity space. By definition, the top five PCs during the auditory period capture a large amount of variance from this period. However, we found that they captured minimal variance from the movement period (monkey 1: 76.9% vs. 3.3%; monkey 2: 88.8% vs. 10.9%, auditory vs. movement period variances, respectively, for PC1 to 5 combined; Fig. 3d). Similarly, the PCs calculated for the movement period captured a large proportion of movement period variance, but a minimum of auditory period variance (monkey 1: 4.8% vs. 86.5%; monkey 2: 12.0% vs. 92.8%, auditory-vs. movement-period variances, respectively, for PC1 to 5; Fig. 3d). This result supports the hypothesis that the auditory and movement periods use close to orthogonal subspaces to perform their computations.

We also calculated an alignment index to quantify the subspace overlap between the activities of both signals. Here, an index of zero means no alignment and one full alignment. The alignment index was 0.08 and 0.23 for monkeys 1 and 2, respectively, significantly lower than an alignment index calculated in a random neural sampling distribution (p < 0.05 for both monkeys). This result indicates that the population rearranges its activity for auditory and movement periods in a nearly orthogonal manner. Therefore, since the neuronal population’s activity behaves in an almost orthogonal fashion in the auditory and movement periods, it is possible to implement a linear decoder to generate two orthogonal subspaces in order to explain the neuronal variability population in both periods. Thus, we applied an algorithm (Elsayed et al., 2016) to find two sets of bases that identify principal components of each period in an orthogonal manner (Figs. 3e and f). The top four PCs from the auditory subspace captured 63.7% and 75.8% of the auditory variance in monkeys 1 and 2, respectively. The top four PCs from the movement subspace captured 76.7% and 82.0% of the movement variance for monkeys 1 and 2, respectively. Plotting one against the other revealed that the T and nT categories separate and move in opposite directions during the auditory period. Meanwhile, T elicits a broad trajectory during the movement period while nT remains close to the origin (Figs. 3g and h). There is a striking similarity between the monkeys’ results, indicating that the appearance of these different population organizations at different times in the task is not monkey-dependent.

### Linear stimulus changes modulate behavior and neural activity

Gradually morphing sounds into one another (Fig. 4a) may reveal the degree to which the neuronal activity changes as a function of the physical dimensions of the stimuli or reflect the behavioral outcome computations. By fitting sigmoidal functions to the monkeys’ behavior (Fig. 4b; Table 1) and the neuronal responses to the morphings (Figs. 4c-d) it is possible to observe the relationship between the neuronal activity and behavior. The neuronal responses of an example neuron (Fig. 3c) identified T at morphing values between 60% and 100%, and nT at morphing values between 0% and 40%. Remarkably, a stimulus at 50% produced a neuronal response between the two groups. However, there was a gradual change during the auditory period, whereas the responses transition more abruptly in the movement period. Figure 4d shows this more precisely. The neurometric and psychometric functions are compared (gray and black lines, respectively) during the auditory, delay, and movement periods. During the latter periods of the task, the neurometric curves of all recorded neurons became more similar to the psychometric curve, i.e. with steeper slopes around the 50% midpoint (Fig. 4e). The overall ratios of the neurometric/psychometric midpoint slopes increased progressively from the auditory, to the delay, to the movement periods: 0.47, 0.60, and 0.93 for monkey 1, and 0.38, 0.69, and 0.92 for monkey 2. Furthermore, figure 4f shows increasing Spearman correlations between neurometric and psychometric curves for all recorded neurons as the task progresses. These results imply that SMA neurons have more information about sensory stimuli during the auditory period and more categorical information (thus becoming more similar to the monkeys’ behavior) during the movement period.

**Table 1.** Overall fitting coefficients of psychometric curves.

| | non-Target | Target | Number of sessions | Number of neurons | Slope | $\theta$ | $\beta_0$ | $\beta_1$ | $\beta_2$ | $R^2$ |
| --- | --- | --- | --- | --- | --- | --- | --- | --- | --- | --- |
| Monkey 1 |  |  |  |  |  |  |  |  |  |  |
|  | grunt | coo | 7 | 24 | 2.0 | 35.2 | 34.9 | 0.6 | 41.7 | 0.80 |
|  | grunt | moo | 6 | 36 | 1.9 | 60.3 | 51.3 | 0.5 | 51.0 | 0.91 |
|  | grunt | w arble | 12 | 57 | 2.3 | 45.6 | 47.3 | 0.5 | 51.7 | 0.96 |
|  | grunt | ['pw ɛΓ.ta] | 10 | 47 | 1.9 | 59.6 | 52.2 | 0.4 | 55.9 | 0.93 |
|  | grunt | [ko.'mi.ð̥a] | 8 | 38 | 1.8 | 60.2 | 56.2 | 0.4 | 52.5 | 0.93 |
|  | [si] | moo | 2 | 10 | 1.6 | 46.7 | 46.6 | 0.4 | 45.5 | 0.94 |
| Monkey 2 |  |  |  |  |  |  |  |  |  |  |
|  | grunt | coo | 13 | 91 | 1.6 | 52.9 | 52.0 | 0.3 | 59.4 | 0.96 |
|  | grunt | moo | 3 | 19 | 2.6 | 45.6 | 52.3 | 0.6 | 51.9 | 0.96 |
|  | grunt | w arble | 3 | 24 | 1.5 | 51.6 | 51.7 | 0.3 | 56.6 | 0.94 |
|  | ['xaw .la] | ['pw ɛΓ.ta] | 8 | 52 | 2.3 | 36.6 | 46.2 | 0.5 | 52.2 | 0.95 |

**Figure 4.**
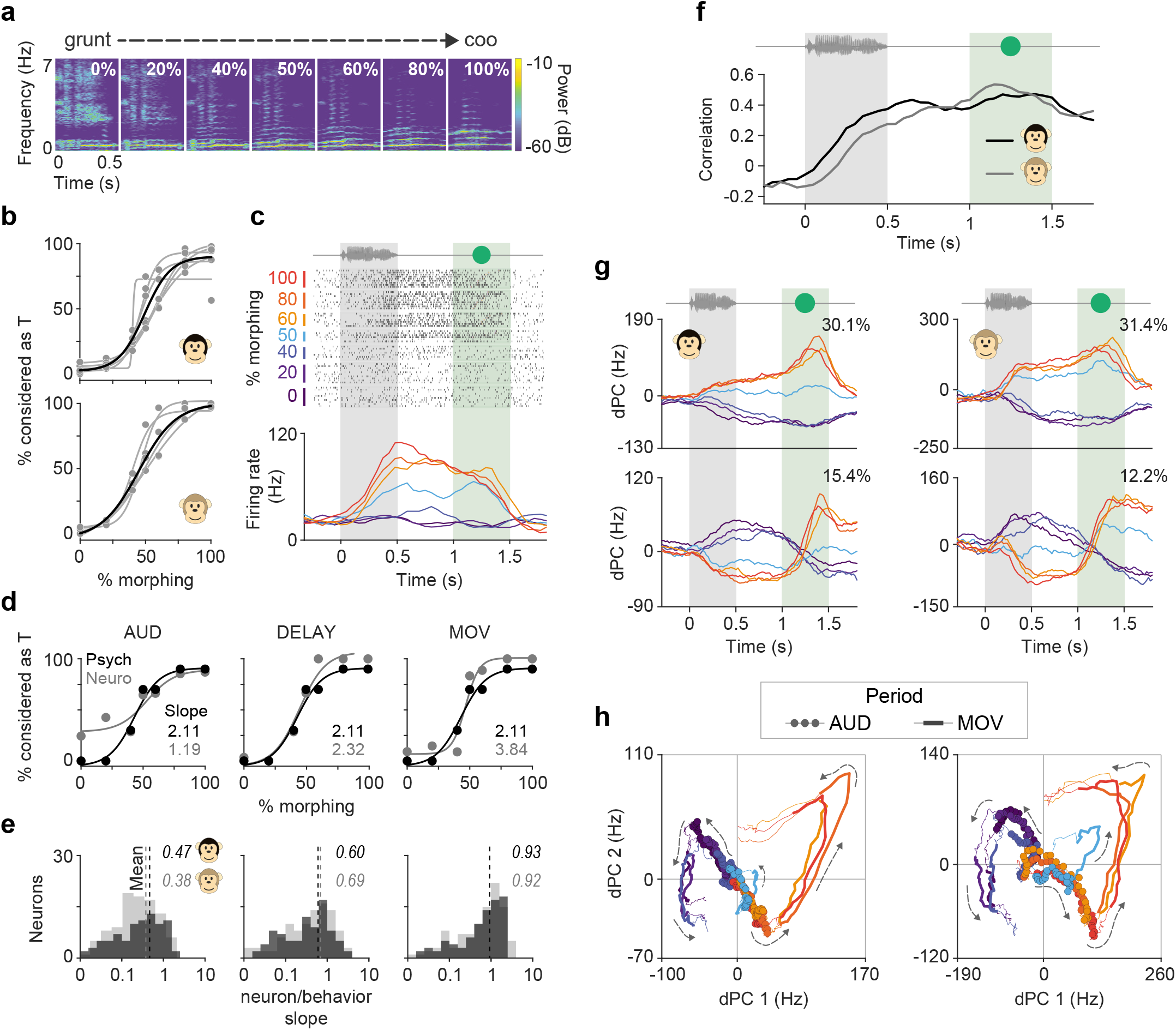
Neural and behavioral responses to morphing sounds. **a)** Spectrogram examples of gradually morphing sounds from grunt (nT) to coo (T). Every morphing set comprises seven morphed sounds. The percentage of T contribution in each sound is at the top of each spectrogram. **b)** Psychometric functions of monkeys (1: top; 2: bottom) as a function of percentage of morphing. Each line is a sigmoidal fit to the behavioral performance. The black line shows the mean performance for all morphings. **c)** Raster (top) and PSTHs (bottom) of a representative SMA neuron during the morphing set (grunt-coo) for monkey 2. **d)** Psychometric (black) and neurometric (gray) comparison in three periods of the neuron in c: auditory, delay, and movement. Each plot shows the percentage of times that behavior or neuronal response considered each morph as T. **e)** Slope ratios (neurometric/psychometric) of all neurons in three periods: auditory, delay, and movement. Monkeys 1: black, 2: gray. Dashed lines and numbers represent means of the distributions (monkeys 1: black; 2: grey). Note that the neurometric curves’ slopes are smaller in the auditory period and become more similar to the psychometric curves during the movement period. **f)** Time-dependent Spearman correlations between neurometric and psychometric data points for both monkeys. **g)** Projection of the first two stimulus dPCs in both monkeys (1: left; 2: right). Colors correspond to a particular percentage of morphing (color code in c). **h)** Two-dimensional projection with the first two dPCs of the population activity for both monkeys (1: left; 2: right)—color code as in g. Circles and thick lines show auditory and movement periods, respectively. Arrows indicate time flow.

Demixed PCA revealed that, in response to morphing stimuli, the population activity followed a pattern similar to that of the single neurons, distinguishing between T and nT and with an in-between response at 50% morphing (Fig. 4g). Divergence started during the auditory period, followed by a separation to two distinct attractors during the movement period (Figs. 4g-h). The two-dimensional dPC projections of the population activity (Fig. 4H) show how neuronal responses to T and nT sounds form two groups and move towards one of two attractors. The transition from one attractor to the other occurred abruptly, further evidencing the SMA’s role in the categorical coding of naturalistic sounds.

### The non-alert condition and errors also point to a role played by SMA in acoustic decisions

To further distinguish between sensory and categorical coding, we projected the neural activity during the errors over the categorical dPCs calculated in correct responses. We found an inversion of activity resembling that of the correct trials (Fig. 5a). That is, in error trials the population activity in SMA reflected the behavioral decision made, rather than the sensory stimulus presented. Moreover, such population activity suggests that when a signal separation during the auditory period does not occur, monkeys make errors. We also recorded the neural activity during sets of passive listening, i.e. when the monkeys listened but did not perform the task because we withdrew the lever and the visual go-cues. Figure 5b shows that during these sets, SMA ceased coding for auditory or categorical decisions. This result indicates that SMA is only active when the monkeys engage in the task, further supporting the role of SMA involvement during acoustic-driven decision making.

**Figure 5.**
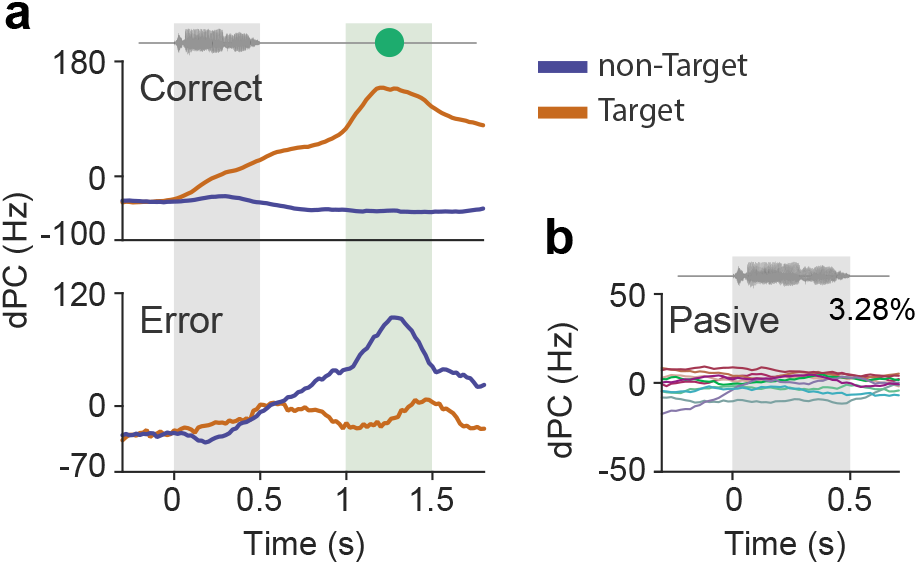
Error trials and the non-alert condition point to a participation of SMA in acoustic decisions. **a)** The categorical dPCs calculated from correct trials (top) were used to project the population neural activity in error trials (bottom) in monkey 1. Note that when the dPCs do not separate during the auditory period, the monkeys make errors. **b)** Projection of first dPC over the neural population during passive trials shows no modulations.

## Discussion

To study whether the SMA codes for acoustic decisions, we trained two monkeys to categorize naturalistic sounds. Observing the time courses of the neural responses, analyzed in different ways, we present here several lines of evidence supporting the concept that SMA integrates sensory information during the auditory period, in support of categorical representations during the movement period: A) Demixed PCA, in which the main dPC began to separate in the auditory period, remained constant during the delay, and then rose sharply during the movement period. B) The two different organizational states were indicative of two different computations being performed in each, during the auditory and movement periods. C) Single neuron recordings during morphing sets showed coding-shifts from sensory to categorical representations as the task progressed. D) The activity of the neuronal population diverged to two distinct attractors, i.e. T and nT. E) Trials in which the monkeys made errors suggested that a separation in the neural population activity seems to be needed during the auditory period; additionally during the movement period the neuronal population with false alarms, had the same activity as during hits.

In previous work with monkeys trained to solve a delayed match-to-sample acoustic task, Scott et al. (2012) reported that the monkeys performed poorly in such tasks. In our study, although the monkeys’ training was more prolonged than in visual or tactile paradigms (Lemus et al., 2009), the monkeys ended up being highly proficient in the categorization of numerous naturalistic sounds (see also Melchor et al., 2020). Therefore, our task presents multiple advantages for studying the neural basis of listening. However, even when the task comprised a great diversity of complex sounds, the monkeys learned to include them in just two categories, i.e. T and nT. Therefore, we did not explore each sound’s semantics, but probably only synonyms, i.e. that multiple sounds mean T, while others mean nT. Moreover, since the monkeys expressed their decisions by releasing a lever or holding it down waiting for a T, one could argue that the monkeys only had to recognize one of the two categories of sounds. Nevertheless, this argument is not plausible since SMA neurons, like the monkeys, acknowledged both categories.

As expected for a premotor area (Nachev et al., 2008; Lara et al., 2018; Shima & Tanji, 2000), during the task’s movement period, single neurons and population responses coded for the monkeys’ dichotomous decisions in the task. Furthermore, dPCA showed that the task’s categorical component captured most of the variance, as opposed to other commonly observed results with this method, in which the condition-independent components are often those with the most weight (Romo et al., 1999; Kobak et al., 2016). Interestingly, the second principal component (Fig. 3a) rises before each auditory stimulus in a predictable fashion, apparently consistent with models of accumulation of information (Leon & Shadlen, 1999). Despite the fact that such a phenomenon, similar to a hazard function of the probability of appearance of a T, was not previously described for auditory processing in SMA, it is consistent with its role in premotor processing (Shima & Tanji, 2000; Nachev et al., 2008; Zimnik et al., 2019). Furthermore, this is the first description of how SMA neuronal activity correlates with decisions based on active listening to naturalistic sounds.

Interestingly, a small group of neurons coded for sounds during auditory periods, as predicted by imaging studies that suggested that the premotor cortex is activated by auditory stimuli (Lima et al., 2016; Vergara et al., 2016; Archakov et al., 2020). Nevertheless, to our knowledge, our results constitute the first electrophysiological evidence of SMA coding natural sounds. Previous fMRI studies using acoustic morphings (Jiang et al., 2018) suggested a two-step category-learning process. First, a perceptual stage in the auditory cortex, and later a categorization stage in the lateral prefrontal cortex. While our current data do not contradict those findings, it adds further nuance by providing evidence that there is a separate computation that processes the physical features of sensory stimuli in premotor cortex. Elsayed et al. (2016) showed that a population of neurons in motor cortex acts as two coordinated circuits of preparatory and movement computations. When we applied their analytical method, we, too, found that the activity of a single neural population in SMA occupies orthogonal subspaces during the auditory and movement periods, implying different computations in each period. Therefore, those two periods’ computations require no more than a single population of neurons organizing itself into two different circuits.

We presented the monkeys with sounds that morphed gradually into one another, in order to determine how SMA responds to linearly changing physical stimuli. Measuring the slope at the midpoint of the sigmoidal curve of the neurometric and psychometric curves, we found that during the auditory period, the neurometric curves have smaller slopes than the psychometric curve, implying that at this moment of the task the neural response is more gradual, and varies more linearly with the changing sensory stimulus. During the movement period, both the neurometric and the psychometric curves had similar slopes, indicating that the neural responses are as categorical as the behavioral choices the animals make. From a population perspective, dPCA shows that sounds are grouped into two clear categories, with the dPC of the 50% response in between these groups. Furthermore, as revealed by the first and second principal components, the population response evolves towards two distinct attractors corresponding to the T and nT categories. The transition from one of these attractors to the other is abrupt rather than gradual, which is a hallmark of categorical coding (Wills et al., 2005; Freedman et al., 2001).

Taken together, our results show how the SMA participates in naturalistic sound-driven decision making by integrating acoustic information to produce decision signals and motor commands. Future work in other cortical areas that lie upstream from the SMA may provide additional information with regard to how this categorization occurs.

## Methods

### Ethics statement

Animals were handled following the Official Mexican Standard for the Care and Use of Laboratory Animals (NOM-062-ZOO-1999) and the study was approved by the Internal Committee for the Use and Care of Laboratory Animals of the Institute of Cell Physiology, UNAM (CICUAL; LLS80-16).

### Animals and experimental setup

Two adult rhesus macaques (*Macaca mulatta*; monkey 1: a 10-year-old male weighing 13 kg; Monkey 2: a 10-year-old female weighing 7 kg) participated in the experiments. Training and recording sessions took place in a soundproof booth. Each monkey sat on a primate chair with its head fixed at a distance of 60 cm from a 21 color LCD monitor (1920 x 1080 resolution, 60 Hz refresh rate). A Yamaha MSP5 speaker (0.05-40 kHz frequency range) was situated fifteen centimeters above and behind the monitor delivering sounds at ~ 65 dB SPL (measured at the monkeys’ ear level). Additionally, a Logitech^®^ Z120 speaker was placed directly below the Yamaha speaker to generate background white noise at ~ 55 dB SPL. Finally, a metal lever placed at the monkeys’ waists captured their responses.

### Task

We trained two rhesus macaques to perform an acoustic recognition task. During the task, the monkeys categorized sounds as either Target (T) or non-target (nT). In a trial, a gray circle of 3° aperture appeared at the center of the screen. Then, a sequence of one to three sounds arose. Each 0.5 s sound continued with a 0.5 s delay and then a 0.5 s green visual cue that replaced the initial gray circle. The cue indicated that the monkey should release the lever if the sound consisted of a T. The monkeys received a reward for releasing the lever in a time window of 0.7 s from the start of the T’s cue. However, after an nT, the monkey kept the lever pressed, waiting for a T. If the monkey released the lever before a T, the trial was classified as a *false alarm*, and a new one commenced. We required that the monkeys’ performance in this task remain above an 80% hit rate before the electrophysiological recordings were made. The programming task was done in LabVIEW 2014 (64-bit SP1, National Instruments^^®^^).

### Auditory stimuli

The sounds were recorded in our laboratory or downloaded from open access online libraries (https://freesound.org/). The monkeys learned more than 37 sounds from 4 different categories (Melchor el al., 2020): conspecific vocalizations, heterospecific vocalizations, words, and artificial sounds. However, for statistical repetitions the recordings were made with only 14 sounds (T = 7; nT = 7): conspecific vocalizations (T = 2; nT = 1), interspecific vocalizations (T = 1; nT = 1) and words (T = 2; nT = 3). The sounds’ sample rate was 44.1 kHz; their amplitudes were normalized at −10dB SPL (RMS), and their durations were shortened or lengthened to 0.5 s (Adobe Audition^®^).

### Morphing sounds

We created sets of morphing sounds that change from an nT to a T in a continuum sequence (STRAIGHT software; Kawahara et al., 1999). This procedure consists of a linear interpolation of five spectro-temporal characteristics of a pair of sounds, i.e. fundamental frequencies, formants, duration, spectro-temporal density, and aperiodicity. The resulting sounds contained some percentage of stimulus A and a complementary percentage of B. Thus, each set consisted of seven morphed sounds (i.e., 0 20 40 50 60 80 and 100 % T). The monkeys trained in various sets; however, monkeys 1 and 2 executed 6 and 4 sets during recordings.

### Passive condition

In this condition, the sounds appeared in a continuous clip separated by 0.3 s interlapses, and with no other cue or visual stimulus. The monkeys did not perform since the lever was not present. However, the monkeys received a reward randomly every 15-30 s.

### Neural recordings

Extracellular recordings of neurons proceeded from an array of five independently movable microelectrodes (1-3 MΩ; Thomas Recording^®^) inserted at different SMA locations in each session. We positioned 20 mm diameter recording chambers according to the Paxinos brain atlas coordinates (Paxinos et al., 2000) and structural magnetic resonance imaging of both monkeys (interaural 27 mm, left hemisphere: lateral 6 mm). The sampled extracellular membrane potentials were 40 kHz (Plexon^®^). Isolation of individual neurons was performed manually using Plexon offline sorter software (Plexon^®^; monkey 1) and automatically using MountainSort (Chung et al., 2017; monkey 2).

### Data analysis

#### Individual neural coding

To evaluate whether SMA neurons coded for T and nT categories or the identity of particular sounds, we performed F-statistics in a one-way ANOVA of the T vs. nT dependent activity and individual sounds. We aligned the neuronal activity to the start of sounds and calculated the mean firing rate in 200-ms windows sliding in steps of 20 ms. To avoid biases in the F-statistic due to comparing classes of different numbers of trials, we equalized the number of trials using the lowest common number of trials in all classes. We repeated the analysis 1000 times to create an F-statistic distribution. To determine the time bins where the F-statistic was significant, we compared the actual distribution against an F-statistic random distribution obtained by shuffling the stimulus labels 1000 times. A neuron was significant if the 95% confidence intervals of the actual and random distributions did not overlap in at least one time bin.

#### Population analysis with PCA

We used principal component analysis (PCA) to describe the SMA’s neuronal population dynamics during the task. We calculated the neurons’ firing rate in blocks of correct trials of 1, 2, or 3 stimuli. We calculated the firing rate using a window of 200 ms moving in steps of 20 ms. Subsequently, we concatenated the firing rate of these three types of trials to obtain a matrix 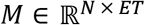 of N number of neurons, E types of trials (with 1, 2, or 3 stimuli), and T total time points. Next, we used the demeaned firing rate of matrix M to calculate the principal components (PCs), considering each row as a variable.

#### Population analysis with demixed PCA

We performed demixed principal component analysis (dPCA; Kobak et al., 2016) in order to observe the effects of the task parameters on the population responses. During the supervised part, dPCA decomposes the neural activity for each variable using the covariance matrices of different marginalizations. The unsupervised analysis of dPCA was similar to that of the PCA, for each marginalization matrix. We marginalized the population activity for T and nT and time. We studied the neuronal activity within times from the starts of sounds to the monkeys’ response, in 200 ms windows sliding every 20 ms. The dPCA algorithm separates the matrices from the decoder (D) and the encoder (F) for each parameter of the task by minimizing the loss function:

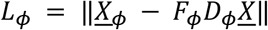

Where *ϕ* denotes the marginalization for each parameter and 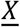 is the mean-centered population matrix (the average of each neuron’s activity is 0). Each component can be ordered based on the amount of variance explained. The axes obtained by the decoder and encoder allow reduction of the data in a few components that capture the most significant variance of each parameter of the task.

We implemented a method to determine the time bins with statistically correct decoding (Kobak et al., 2016). First, we separated the data into training and testing sets. We constructed the testing set by randomly separating one trial from each category (T and nT). In contrast, the training set was the average of the rest of the trials in each category. We performed dPCA from the training set to obtain the decoding axes. Then we projected the testing set on these axes and classified them according to the closest category mean (T or nT). We repeated this procedure 1000 times and measured the proportion of correct classification. We used 100 shuffles to calculate the classification accuracies expected by chance. The shuffled distribution was computed by choosing random trials between categories for each neuron. We ran the same method described above and calculated the proportion of correct classifications for each iteration. Then we compared the proportion of correct classifications between the real data vs. the shuffled data. The time bins where the real classification exceeded the shuffled classification distribution were considered statistically different from chance.

#### Identifying different subspaces in auditory and movement periods

To compare the population states during the auditory and movement periods, we performed four procedures to describe the transition between one to other state and a new algorithm developed by Elsayed et al. (2016) to calculate optimally two orthogonal subspaces, where each one captures auditory and movement activity. 1) To compare the population covariation between auditory and movement periods, we computed the correlation between all neurons in each period. We created two matrices for each period: *A ∈ R^N×CT^* and *M ∈ R^N×CT^* where N is the number of neurons, C is the number of stimuli and T is the time. The correlation matrix of each period was obtained computing the Pearson correlation between the rows of each matrix (A and M). 2) To quantify the subspace overlapping between subspaces of both periods, we computed a PCA over *A ∈ R^N × CT^* and *M ∈ R^N × CT^* to obtain the top ten auditory and movement PCs. First, the firing rate of each period was projected onto the auditory PCs and the percentage of explained variance relative to the total variance of the auditory period was quantified. Then, the same procedure was executed for movement PCs. 3) We calculated an alignment index to measure the amount of auditory period variance captured by the top ten movement PCs normalized by the auditory period variance captured by the top ten auditory PCs. The index range varied from 0 to 1, where 0 indicates the orthogonality between both periods and 1 means a perfect alignment. 4) To rule out that two groups of neurons are activated differentially in the auditory and movement periods we use an epoch-preference index, which evaluates the relative weight of each neuron in both periods. We calculated the firing rate range separately in each period and divided it by the mean of the firing rate of the entire trial. Then the range of each neuron was normalized by the mean of the firing rate of all neurons. A neuron with a positive value indicates a preference for the auditory period over the movement period, a negative value indicates the opposite. A bimodal distribution of the index signifies the existence of two neural subpopulations, one that is more selective for auditory period and other for movement period. Finally, we used a method developed by Elsayed et al. (2016) to derive two orthogonal subspaces, each one capturing auditory and movement activity, respectively. This method maximizes the sum of the variance of each period activity in each period subspace and was calculated with the following loss function:

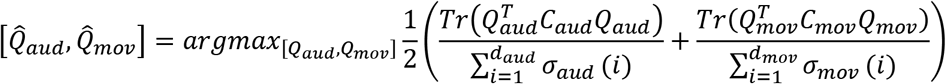

Where *C_aud_* and *C_mov_* are covariance matrices of the neural activity in auditory and movement period, respectively. *σ_aud_*(*i*)_*y*_ *σ_mov_*(*i*) are ith singular value of the *C_aud_* and *C_mov_*, respectively. *Q_aud^y^_ Q_mov_* are the eigenvectors of auditory and movement subspaces. We selected four dimensions for *Q_aud_* and *Q_mov_* due they capture close to 70% variance of each period. This method identifies simultaneously the auditory and movement subspaces while constraining them to be orthogonal.

#### Psychometric and Neurometric Measures

We adjusted hyperbolic tangent functions to the psychophysical and neurometric performance during morphing sets. This function relates the percentage of times in which the monkey (psychometric curve) or the neuron (neurometric curve) recognized a sound like a T as a function of a morphing sound. The function is:

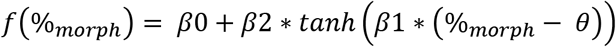

θ and β0 represent the midpoint of the function on the horizontal and vertical axis, respectively, β2 represents the amplitude function from β0, and β1 defines the morphing percentage dependent of behavior, a high value indicates an acute sigmoid. Additionally, we measured a linear slope at the sigmoid’s midpoint to assess the degree of perceptual and neural modulation as a function of the morphing rate. We calculated the maximum and minimum values of the sigmoid’s second derivative to find the central part of the sigmoid. This section corresponds to the behavioral transition in the categorization of morphing stimuli. After obtaining the abscissa’s and the ordinate’s coordinates, we used the following equation to determine the slope:

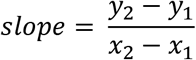

#### Neurometric function

We derived the neurometric functions from each neuron’s correct trials (de Lafuente & Romo, 2005). A neurometric curve represents the probability that an ideal observer reports the stimulus category (T or nT) based solely on the firing rate. For each neuron and time bin, the neurometric curve was the percentage of trials in which the maximum firing rate reached or exceeded an arbitrary threshold. The threshold was selected to maximize the number of correct answers. For the neurometric curves, we used 300 ms windows, sliding every 50 ms. Psychometric and neurometric sensitivity were compared based on sigmoidal slopes. We used a Spearman’s correlation between the raw data from the psychometric and neurometric curves.

## Acknowledgements

We thank Elizabeth Cabrera for valuable comments. We are also grateful for the financial support provided by CONACYT, CB-256767, and Programa de Apoyo a Proyectos de Investigación e Innovación Tecnológica (PAPIIT) IN207919. Special thanks to Gerardo Coello and Ana Escalante of the computing department of the IFC, to Gabriel Pérez Ruelas for technical assistance, and to Mr Patrick Weill for proofreading the manuscript. Isaac Morán Martinez is a doctoral student from Programa de Doctorado en Ciencias Biomédicas, Universidad Nacional Autónoma de México (UNAM) and received fellowship 288348 from CONACYT, México. Data in this work are part of his doctoral dissertation.

## Author Contributions

IM, JM, TF & LL performed experiments. IM, & JPO analyzed data and prepared figures. LL designed the paradigm. TF programmed the task. JPO & LL wrote the paper.

